# Genetic and Physiological Characterization of the Antibacterial Activity of *Bacillus subtilis* subsp. *inaquosorum* Strain T1 Effective Against *pirAB*^*Vp*^-Bearing Vibrio *parahaemolyticus*

**DOI:** 10.1101/2020.08.07.242404

**Authors:** Sarah E. Avery, Susannah P. Ruzbarsky, Amanda M. Hise, Harold J. Schreier

**Affiliations:** Department of Biological Sciences, University of Maryland Baltimore County, Baltimore, Maryland, USA; Department of Marine Biotechnology, Institute of Marine and Environmental Technology, University of Maryland Baltimore County, Baltimore, Maryland, USA

## Abstract

Acute hepatopancreatic necrosis disease (AHPND) is caused by PirAB toxin-producing *Vibrio parahaemolyticus* and has devastated the global shrimp aquaculture industry. One approach for preventing growth of AHPND-producing *Vibrio* spp. is through the application of beneficial bacteria capable of inhibiting these pathogens. In this study we focus on the inhibitory activity of *Bacillus subtilis* subsp. *inaquosorum* strain T1, which hinders *V. parahaemolyticus* growth in co-culture experiments in a density-dependent manner; inhibition was also obtained using cell-free supernatants from T1 stationary phase cultures. Using a *mariner*-based transposon mutagenesis, 17 mutants were identified having complete or partial loss of inhibitory activity. Of those having total activity loss, 13 had insertions within a 42.6 kb DNA region comprising 15 genes whose deduced products were homologous to non-ribosomal polypeptide synthetases (NRPSs), polyketide synthases (PKSs) and related activities, which were mapped as one transcriptional unit. Mutants with partial activity contained insertions in *spo0A* and *oppA*, indicating stationary phase control. Expression of *lacZ* transcriptional fusions to NRPS and PKS genes was negligible during growth and at their highest during early stationary phase. Inactivation of *sigH* resulted in loss of inhibitor activity, indicating a role for σ^H^ in transcription. Disruption of *abrB* resulted in NRPS and PKS gene overexpression during growth as well as enhanced growth inhibition. This is the first study examining expression and control of an NRPS-PKS region unique to the *inaquosorum* subspecies of *B. subtilis* and an understanding of factors involved in T1 inhibitor production will enable its development for use as a potential tool against AHPND *Vibrio* pathogens in shrimp aquaculture.

**IMPORTANCE:** The shrimp aquaculture industry has been impacted by the rise of acute hepatopancreatic necrosis disease (AHPND), resulting in significant financial losses annually. Caused by strains of the bacterial pathogen, *Vibrio parahaemolyticus*, treatment of AHPND involves the use of antibiotics, which leads to a rise in antibiotic resistant strains. An alternative approach is through the application of beneficial microorganisms having inhibitory activities against AHPND-generating pathogens. In this study, we examine the genetic basis for the ability of *Bacillus subtilis* strain T1 to inhibit growth of an AHPND *Vibrio* strain and show that activity is associated with genes having the potential for synthesizing antibacterial compounds. We found that expression of these genes is under stationary phase control and showed that inactivation of a global transition state regulator results in enhancement of inhibitory activity against the AHPND *Vibrio*. Our approach for understanding the factors involved in production *B. subtilis* strain T1 inhibitory activity may allow for development of this strain for use as a potential tool for the prevention of AHPND outbreaks.

## INTRODUCTION

Acute hepatopancreatic necrosis disease (AHPND) in juvenile penaeid shrimp, also known as early mortality syndrome (EMS), first emerged in China in 2009 (1) and has spread to Vietnam, Malaysia, Thailand, Mexico, the Philippines and throughout South America (1) as well as the United States (2). AHPND leads to 100% mortality and has resulted in annual losses for shrimp farming estimated to be over 1 billion US dollars (3). The disease is caused by strains of *Vibrio parahaemolyticus* that carry a plasmid encoding a *Photorhabdus* insect-related binary toxin PirAB, also known as Pir-likeAB and PirVP (4-6). AHPND is lethal in both *Penaeus monodon* and *Litopenaeus vannemei*, the two most cultivated shrimp species in the aquaculture industry (3). Treatment has led to the inevitable increase in resistance to commonly used antibiotics, e.g., oxytetracycline, quinolones and amoxicillin (7, 8). Thus, alternative approaches for preventing and treating this disease are required.

Colonization by AHPND-causing *V. parahaemolyticus* strains creates a major shift in the microbiota of the shrimp gut (9). In healthy shrimp, Rhodobacterales and Rhizobiales along with Planctomycetales are the major contributors to the gut microbiome. Post-infection, the Mycoplasmatales and Vibrionales are the dominant gut inhabitants (10). The practice of draining and disinfecting ponds between shrimp stocks may increase the risk of an AHPND outbreak by removing beneficial bacteria. Manipulating microbial communities using “microbially mature” water has been shown to increase survival of fish larvae over the use of filter-sterilized water (11). Thus, one approach to combating this disease is through the use of beneficial bacteria— probiotics—capable of positively modifying shrimp microbial communities and, at the same time, interfering with growth of pathogens like the AHPND-causing strains. Probiotics are live microorganisms that, when administered in adequate amounts, confer health benefits to the host (12). These benefits occur through a variety of mechanisms, including the production of antimicrobial compounds (12).

A group of bacteria that have received attention for their probiotic potential are the Gram-positive spore-forming *Bacillus* sp. due to the antimicrobial activities of their structurally diverse secondary metabolites including polyketides, aminoglycosides, and nonribosomal peptides such as bacteriocins and lipopeptides (13, 14). Production of these antimicrobial compounds is usually associated with stationary phase physiology, which is associated with adverse changes in their environment. Biosynthetic pathways for these activities are often organized as operons and include nonribosomal peptide synthetases and polyketide synthases, which are modular in design and are arranged in various combinations resulting in the production of different compounds (15, 16). Many act through permeabilization and destruction of the cell membrane and other mechanisms (15).

The present study focuses on *Bacillus subtilis* subsp. *inaquosorum* strain T1, which we have found to possess an inhibitory activity against AHPND-producing *Vibrio parahaemolyticus* strains (17). As a member of the *inaquosorum* subspecies [(17) and Schreier, unpublished], it is similar to the *stercoris* subspecies of *B. subtilis* but differs from *subtilis* and *spizizeni* subspecies due to coding capacity for an uncharacterized nonribosomal peptide synthetase (NRPS)-polyketide synthase (PKS) gene cluster (18). In this study we demonstrate that T1 inhibitory activity is associated with a secreted product and we demonstrate the involvement of the NRPS-PKS-encoding region in this activity. We found that NRPS-PKS gene expression was negligible during mid-exponential growth and at highest during stationary phase and that stationary phase regulators *oppA (spo0K), spo0A*, and *sigH (spo0H)* are involved in their control. Finally, disruption of the global transitional phase regulator *abrB* resulted in derepressed NRPS-PKS expression during exponential phase and enhanced inhibitory activity. Our study examines expression and control of an NRPS-PKS region unique to the subspecies *inaquosorum* of *B. subtilis* and we suggest that our ability to manipulate strain T1 to be an effective inhibitor against an AHPND-producing *Vibrio* strain may be one approach for developing tools for use as probiotics against AHPND in shrimp.

## METHODS

### Bacterial Strains and Culture Media

Bacterial strains used in this study and their sources are listed in Table 1. Strains were grown in Zobell 2216 marine broth (HiMedia Laboratories), tryptic soy broth (Sigma-Aldrich) or agar supplemented with 2% NaCl (TSB2 or TSA2, respectively) and lysogeny broth (LB) agar (19) with antibiotic, when appropriate. SOC and 2XYT have been described (19). Antibiotics were added to media at concentrations of 100 μg spectinomycin (Sp)/ml, 1 μg erythromycin (Em)/ml and 10 µg lincomycin (Ln)/ml. Strain T1g was constructed by transforming strain T1 by electroporation (described below) with DNA from *B. subtilis* strain AR13 (*amyE*::*gfp)* and selecting for spectinomycin-resistance (Sp^R^). Incorporation of *gfp* into *amyE* was confirmed by the loss of amylase activity on starch agar and by an increased size of *amyE* by PCR. Disruption of *amyE* did not affect D4 inhibitory activity as determined by overlay or co-culture assays (see below); T1 and T1g were interchangeable and their choice for usage depended on the experiment. Strains SSh1*(sigH::erm*) and SSa1(*abrB::erm*) were constructed by transforming strain T1 with DNA from *B. subtilis* strains BKE00980 (*sigH::erm*) and BKE00370 (*abrB::erm*), respectively, selecting for Em- and Ln-resistance; insertion of the *erm* cassette was confirmed by PCR. *Vibrio parahaemolyticus* strains D4, isolated from Mexico, and A3, isolated from Viet Nam, were provided by Dr. Kathy Tang-Nelson, University of Arizona and the presence of *pirAB* was confirmed by PCR.

**Table 1.** Bacterial strains used in this study.

| Strain | Description | Source or Reference |
| --- | --- | --- |
| T1 | *Bacillus subtilis* subsp. *inaquosorum* | Epicore Networks (U.S.A.) Inc. |
| SMY | *Bacillus subtilis* subsp. *subtilis* | Laboratory strain |
| T1g | *amyE::rrnA<sub>p</sub>-gfp*, T1 x AR13 DNA, Sp<sup>R</sup> | This study |
| SSh1 | *sigH::erm*, T1 x BKE00980 DNA, Em<sup>R</sup> Ln<sup>R</sup> | This study |
| SSb1 | *abrB::erm*, T1 x BKE00370 DNA, Em<sup>R</sup> Ln<sup>R</sup> | This study |
| AR13 | *amyE::rrnA<sub>p</sub>-gfp* Sp<sup>R</sup> | BGSC |
| BKE00980 | *sigH::erm* | BGSC |
| BKE00370 | *abrB::erm* | BGSC |
| D4 | *Vibrio parahaemolyticus* isolate 13-306, PirAB | (4) |
| A3 | *Vibrio parahaemolyticus* isolate 13-028, PirAB | (4) |
| ATCC 17802 | *Vibrio parahaemolyticus* | James Kaper |
| A1-20 | *orfDΩTnLacJump (orfD::lacZ)*, Sp<sup>R</sup> | This study |
| A2-18 | *oppAΩTnLacJump*, Sp<sup>R</sup> |  |
| A2-30 | *orfBΩTnLacJump*, Sp<sup>R</sup> |  |
| A3-41 | *orfBΩTnLacJump (orfB::lacZ)*, Sp<sup>R</sup> |  |
| A8-11 | *spo0AΩTnLacJump*, Sp<sup>R</sup> |  |
| A10-67 | *orfBΩTnLacJump*, Sp<sup>R</sup> |  |
| A11-79 | *orfCΩTnLacJump (orfC::lacZ)*, Sp<sup>R</sup> |  |
| A13-56 | *orf8ΩTnKRM*, Sp<sup>R</sup> |  |
| A20-86 | *orfAΩTnLacJump (orfA::lacZ)*, Sp<sup>R</sup> |  |
| A23-33 | *orfFΩTnLacJump*, Sp<sup>R</sup> |  |
| S2-30 | *orfBΩTnKRM*, Sp<sup>R</sup> |  |
| S11-14 | *orfBΩTnKRM*, Sp<sup>R</sup> |  |
| S12-29 | *orfBΩTnKRM*, Sp<sup>R</sup> |  |
| S21-28 | *orfCΩTnKRM*, Sp<sup>R</sup> |  |
| S21-46 | *orfAΩTnKRM*, Sp<sup>R</sup> |  |
| S31-22 | *yoaZΩTnKRM*, Sp<sup>R</sup> |  |
| S35-30 | *orfEΩTnKRM*, Sp<sup>R</sup> |  |
| A20-86-A | *abrB::erm*, *orfAΩTnLacJump (orfD::lacZ)*, A20-86 x BKE00370 DNA, Em<sup>R</sup> Ln<sup>R</sup> Sp<sup>R</sup> |  |

### Soft Agar Overlay Assay

To evaluate inhibitory activity, 2.5 µl from an overnight *B. subtilis* culture was spotted onto 2216 agar and incubated at 37°C for 18 hr. In a chemical fume hood, uncovered plates were placed in a pyrex dish along with a reservoir of chloroform (20 to 30 ml); the dish was then covered with plastic wrap to generate a chloroform atmosphere and facilitate cell death. After 30 min, plates were removed, covered, and set at room temperature for 30 min to allow for chloroform evaporation. For each plate, 3 ml of semi-solid 2216 agar (2216 broth with 0.75% Bacto agar) was heated until liquified, cooled to 42°C, inoculated with 10 μl of an overnight *V. parahaemolyticus* culture, and immediately poured over the 2216 agar surface. Plates were incubated at 28°C overnight and examined for zones of clearance.

### Co-Culture Growth Experiments

Growth experiments examining the effect of T1g or mutant A3-41 on D4 growth were done by combining overnight cultures of D4 with T1g or A3-41 at various cell densities in fresh liquid 2216 broth and monitoring D4 by quantitative PCR (qPCR) (see below). The qPCR target to assess densities of T1g, mutant A3-41, and strain D4 were *gfp, amyE*, and *toxR*, respectively, and primers are listed in Table 2. Inocula for co-cultures were based on colony forming units/ml (CFU/ml) for each strain in 2216 broth, which ranged from 2.0 x 10^10^ to 3.0 x 10^10^ CFU/ml for D4 and 2.0 x 10^9^ to 3.0 x 10^9^ CFU/ml for strains T1g and A3-41. Overnight cultures of T1g, A3-41, and D4 were prepared in 2216 broth and were used to inoculate 50 ml of 2216; overnight cultures of T1g and A3-41 were rinsed and suspended in fresh 2216 prior to their use as inoculants. Initial cell density for D4 was 2.0 x 10^4^ CFU/ml and initial cell densities for T1g and A3-41 ranged from 2.0 x 10^4^ to 2.0 x 10^6^ CFU/ml to generate T1g or A3-41 to D4 ratios of 1:1, 10:1, and 100:1, respectively. Cultures were grown at 28°C and 240 RPM for 24 hr. At 3 hr after inoculation, 10 ml of each culture was centrifuged at 4°C, 4,000 x g for 10 min; at 24 hr after inoculation, 0.5 ml of each culture was centrifuged at 4°C, 10,000 x g for 5 min. To assist in the recovery of low-density D4 cultures at the 3 hr time point, autoclaved *Aeromonas hydrophila* was added to each sample to a final concentration of 10^5^ CFU/ml prior to centrifugation. Extraction of DNA from each sample was done using the Wizard Genomic DNA Purification Kit (Promega) following the manufacturer’s specifications.

**Table 2.**
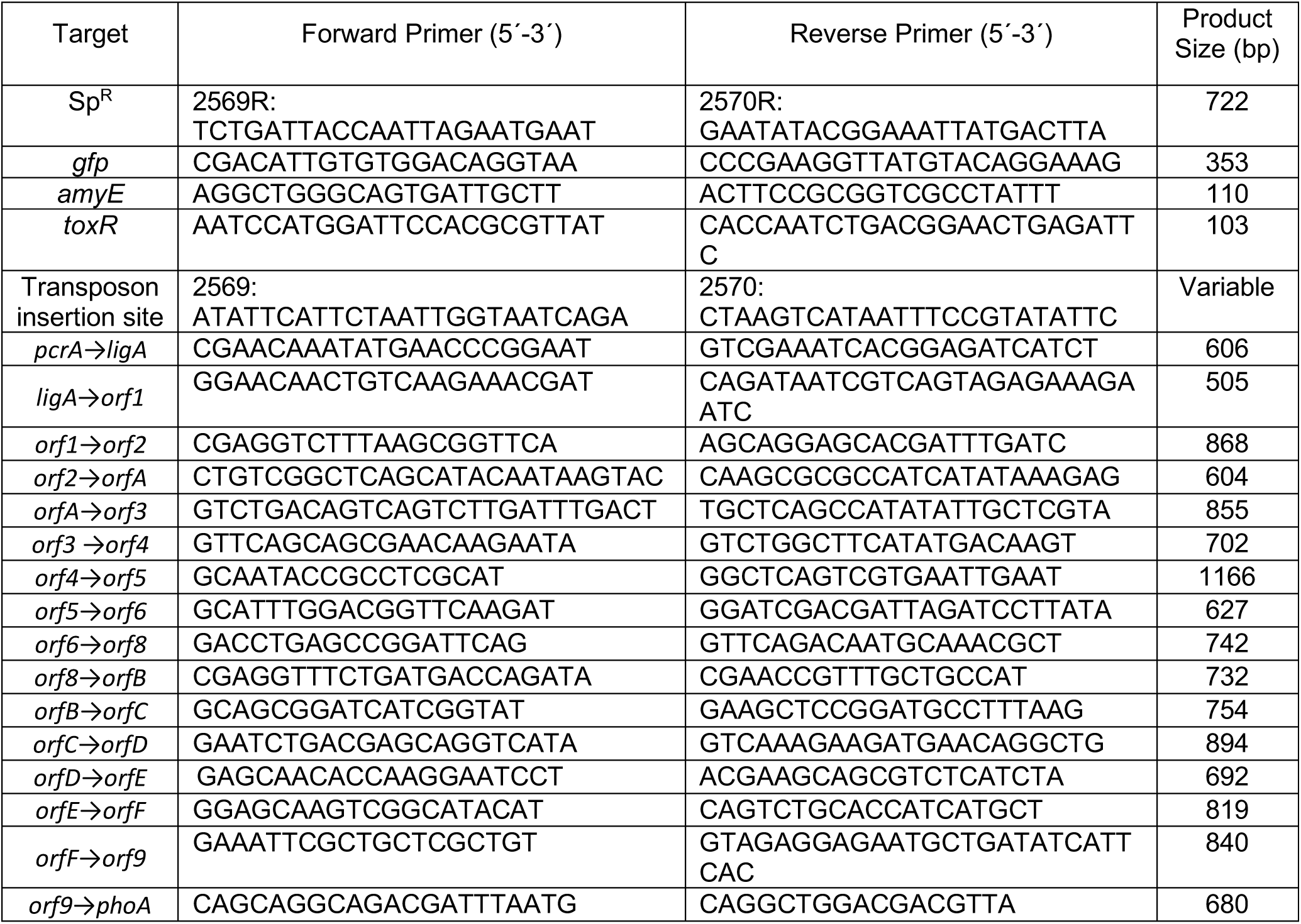
Primers used in this study.

### Cell-free culture supernatant experiments

Overnight cultures grown in 2216 broth were centrifuged at 4°C, 5,000 x g for 10 min and supernatant fractions were passed through a 0.2 μm filter. Filtered supernatants were then added to fresh 2216 medium in 50 ml sidearm flasks to a final concentration of 50% in a total volume of 12 ml. The control flask received 12 ml of fresh 2216 broth. Each culture was then inoculated with D4 at a concentration of 2.0 x 10^4^ CFU/ml and cultures were grown at 28°C and 240 RPM, monitoring growth using a Klett-Summerson Colorimeter with a Wratten 54 filter. Two 1 ml samples were taken from each flask at 5.5 hr after inoculation, followed by DNA extraction as described above.

### qPCR analysis

qPCR was performed in 10 μl reactions containing 5 μl of PefeCTa SYBR Green Fastmix (Quanta), 3.5 μl PCR-certified water, 0.25 μl of 1/10 diluted forward and reverse primers (0.5 μM) each, and 1 μl of the sample to be quantified. All qPCR reactions were performed using an Applied Biosystems 7500 Fast Real-Time PCR machine. PCR-certified water was used for the no template control to monitor for contamination. Primers for qPCR are listed in Table 2. The DNA template for standard curves was prepared by PCR performed in 50 μl reactions containing 25 μl Taq PCR Mastermix (Qiagen), 19 μl PCR-certified water, 2 μl of 1/10 diluted forward and reverse primer (0.5 μM) for *toxR, gfp* and *amyE* (Table 2), and 2 μl of DNA template. Chromosomal DNA from D4, T1g, and T1 were used for the DNA templates for *toxR, gfp*, and amyE, respectively. PCR products were then purified using the MinElute PCR Purification Kit (Qiagen) according to the manufacturer’s specifications and concentration (ng/ml) was determined using a Qubit Fluorometer (ThermoFischer).

### Plasmids

Mariner-derived *himar1* delivery vectors pDP384 (*TnKRMspec amp mls mariner-Himar1ori*(TS)_*Bs*_ and pEP4 (*TnLacJump spec amp mls mariner-Himar1ori*(TS)_*Bs*_) (20) were used for transposon mutagenesis of strain T1 and were provided by Dr. Daniel Kearns, Indiana University. The plasmids harbor a temperature-sensitive *B. subtilis* origin of replication and erythromycin-resistance (Em^R^) located outside of transposon sequences and Sp^R^ contained within transposon sequences. Growth at the restrictive temperature and selection for Sp^R^ resulted in the identification of cells containing chromosomal insertions. Plasmid pEP4 differs from pDP384 by the presence of a promoter-less *lacZ* gene within transposon sequences, which is expressed when inserted downstream of an active transcription start-site (20). Plasmids were purified using a Wizard DNA Clean-Up Kit (Promega) according to manufacturer’s standards.

### Electroporation of T1 with Delivery Vectors and Mutagenesis

T1 was prepared for electroporation by growing in 250 ml of 2xYT to OD_600_=0.8 at 37°C, 240 RPM, washing cells three times in ice cold 10% glycerol and suspending in 1 ml ice cold 10% glycerol, storing at -80°C in 200 µl aliquots. For electroporation, one aliquot was mixed with purified plasmid (∼7 μg/ml) then incubated on ice for 5 min. The cells were then transferred to a 2 mm electroporation cuvette and electroporation was done using the “StA” program of a Bio-Rad MicroPulser electroporation apparatus (∼1.8 kV for 2.5 msec). After electroporation, 0.5 ml SOC was added and cells were incubated at 28°C with aeration for 2 hr, followed by plating onto LB+Sp+Em agar. The presence of the transposase gene in several Sp^R^ Em^R^ transformants was confirmed by PCR and one transformant was selected for mutagenesis. After growth in LB+Sp for 18 hr at 42°C, cells were plated onto LB+Sp agar at a dilution of 10^−6^. Approximately 3,000 transposants were screened for decreased or lost D4 inhibitory activity by the presence of reduced (relative to a wild-type control) or absent clearance zones, respectively, using the overlay assay substituting TSA2 for 2216 agar, as described above.

### Mutant Characterization and Identification of Transposon Insertion Site

Confirmation that Sp^R^ was due to transposon insertion was done by PCR using primer set 2569R/2570R (Table 2) followed by visualization via agarose gel-electrophoresis. PCR was carried out in 25 μl reactions with Qiagen *Taq* polymerase using a Bio-Rad S1000 Thermal Cycle for 3 min at 94°C followed by 30 cycles of 1 min at 94°C, 1 min at 55°C, 1 min at 72°C, and a final step at 72°C for 10 min. To ensure that a mutant had only one insertion, chromosomal DNA was prepared from that mutant and re-transformed into strain T1 by electroporation, selecting for Sp^R^. Insertion site identification was determined by amplification of the transposon and adjacent DNA using an inverse PCR strategy as follows. One μg of chromosomal DNA from T1 mutants was digested with *Taq*I for 2 hr at 37°C in 20 μl and one μl from this reaction was then ligated using the T4 DNA Rapid Ligation Kit (Thermo Scientific) for 5 min at room temperature according to manufacturer’s specifications. The ligation mixture was then used in a PCR reaction with primers 2569/2570 (Table 2) and Phusion DNA polymerase (New England BioLabs). PCR was carried out in 50 μl reactions for 3 min at 98°C followed by 30 cycles of 1 min at 98°C, 1 min at 55°C and 1 min at 72°C, and a final step at 72°C for 10 min. PCR products were purified using the Wizard PCR Preps DNA Purification System (Promega) and DNA sequencing was performed using primer 2569, which anneals to transposon sequences adjacent to the insertion site. Comparison of DNA sequences interrupted by the transposon to database sequences was done by BLAST (21) and BLASTp (22).

### Isolation of RNA

Overnight cultures of T1 grown in 2216 broth were used to inoculate 10 ml of 2216 broth and at ∼1-2 hour before (T_-1_) and after reaching stationary phase (T_1_) 1 ml samples were removed and centrifuged at 10,000 rpm, 4°C, for 5 min, discarding the supernatant fraction. Cell pellets were suspended in 0.3 ml a solution of 10 mM pH 8.0 Tris-HCl and 10 mg lysozyme/ml (Sigma-Aldrich) and incubated at 37°C for 1 hr. TRIzol (Thermo Fisher Scientific) (1 ml) was added followed by 0.2 ml cold chloroform and incubation was continued at room temperature for 2-3 min, then centrifuged at 13,000 rpm, 4°C, for 20 min. After centrifugation, 0.4 ml of the resulting aqueous phase was added to 0.5 ml isopropanol and incubated at room temperature for 10 min to precipitate RNA. The precipitant was washed with 75% ethanol, dried, and suspended in 50 µl DEPC-treated water. RNA quality was assessed by ethidium bromide-agarose gel electrophoresis.

### RT-PCR for transcription mapping experiments

T1 RNA was treated with RNase-free DNaseI (Thermo Fisher Scientific) prior to being used for reverse transcriptase (RT)-PCR to eliminate residual genomic DNA; 50-100 ng RNA was used for each reaction. RT-PCR was carried out using gene-specific primers (Table 2) at a final concentration of 0.5 µM, using the SuperScript IV One-Step RT-PCR System (Thermo Fisher Scientific). Reactions were performed using a reverse transcription step at 50°C for 10 min, a 2-min RT inactivation step at 98°C followed by 25-30 cycles at 98°C for 10 sec, 56°C for 10 sec, and 30 sec/kb at 72°C, with a final extension at 72°C for 5 min. Each primer set (Table 2) was tested with T1 chromosomal DNA as template using *Taq* polymerase.

### ß-Galactosidase assays

Overnight cultures were used to inoculate 2216 medium in sidearm flasks and cultures were grown at 37°C, 240 rpm, monitoring cell density at an OD_600_. Cells (1 ml) were harvested at mid-exponential (T_-1_), at the onset of stationary (T_0_), and 1-2 hours into stationary phase (T_1_), centrifuging at 10,000 x g, 4°C, 5 min., and suspending cell pellets in 1 ml modified Z buffer (23). Sodium dodecylsulfate was added (final concentration 0.05%), vortexed, then incubated at 37°C, 5 min, after which 0.2 ml of o-nitrophenyl-ß-galactoside (4 mg/ml) was added, mixed, and incubated at 37°C for an additional 30 to 60 min. The reaction was stopped by the addition of 0.25 ml 2M sodium carbonate, vortexed, and placed on ice. After centrifugation at 10,000 x g at room temperature for 10 min., the OD at 420 nm was measured. Specific activity is given as nmoles/min/OD_600_.

### Statistical analysis

Analysis of the data was performed using one-way analysis of variance (ANOVA) with the significance level of 0.01 (p < .01).

## RESULTS

### Assessing T1 inhibitory activity

Using the soft-agar overlay assay, T1 was shown to inhibit growth of three strains of *Vibrio parahaemolyticus*— AHPND-causing strains D4 (Fig 1A), isolated from Mexico, and A3 (Fig. 1B), isolated from Viet Nam, as well as a non-AHPND strain, ATCC 17802 (Fig. 1C, left)—which is evident by the zone of clearance around the T1 colony. Because chloroform is toxic to T1, the growth inhibition observed for the *Vibrio* strains applied in the soft agar over the T1 colony is a consequence of the accumulation of a diffusible substance produced by T1. In contrast, SMY, which is a *B. subtilis* subsp. *subtilis* strain (24), did not inhibit growth of any of the three strains (Fig. 1, right), highlighting a difference between *inaquosorum* and *subtilis* subspecies. Because strain D4 was more sensitive to T1 than A3, based on the size of the clearance zone, we used D4 as the test strain for subsequent studies.

**Figure 1.**
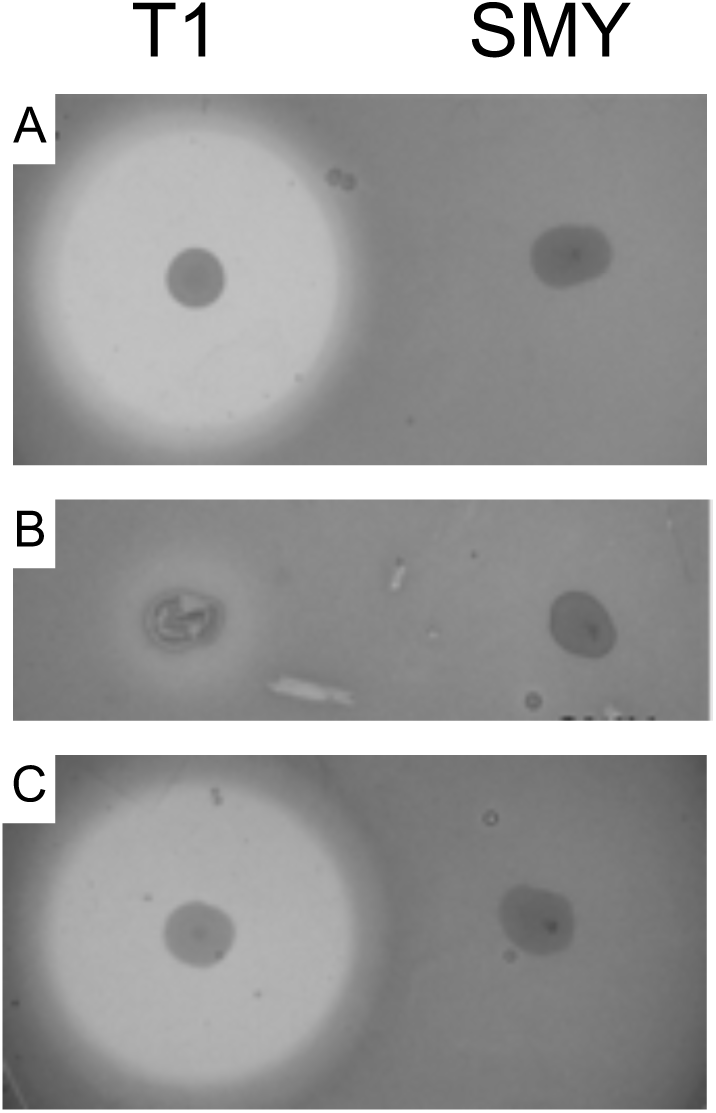
Soft-agar overlay assays for *B. subtilis* strains T1 (left) and SMY (right) against *pirAB*-harboring *V. parahaemolyticus* strains D4 (A) and A3 (B), and a non-*pirAB V. parahaemolyticus* strain (C). Assays were done as described in Methods. A zone of clearance around a colony indicates the absence of growth of the *Vibrio* spp.

The ability of T1 to effect D4 was examined further by co-culturing the two strains and monitoring D4 levels by measuring *toxR* copy number (see Methods). For these growth experiments, overnight cultures of D4 were diluted to 2 x 10^4^ CFU/ml and combined with overnight cultures of strain T1g diluted to final densities of 2 x 10^4^, 2 x 10^5^ and 2 x 10^6^ CFU/ml (1:1, 10:1, and 100:1, T1g to D4, respectively) in 2216 broth, as described in Methods; the control culture did not contain T1g. After 3 hr and 24 hr, samples were collected, and DNA was extracted for qPCR analysis. At 3 hr, *toxR* levels did not vary significantly for any of the treatments (Table 3). At 24 hr, levels increased approx. 10^3^- to 10^4^-fold for control and cultures supplemented with T1g at the 1:1 and 10:1 ratio. On the other hand, D4 *toxR* levels in the 24 hr culture containing T1g at the 100:1 ratio was between 750- to 1250-fold lower compared to control (no T1g) and the other T1-supplemented cultures and increased only approx. 70-fold compared to its 3 hr time point (Table 3). At 24 hr, T1g levels (measuring *gfp* copy number) were not significantly different for all T1-containing cultures, varying between 6×10^7^ ± 1.1×10^7^ copies/ml for the 1:1 culture to 1.6×10^8^ ± 0.6×10^8^ copies/ml for the 100:1 culture. Thus, while T1g levels reached similar cell densities for all three treatments, D4 growth inhibition by T1g could be observed but only when present at a 100-fold excess, suggesting a relationship between inhibitor and culture density as well as an ability for D4 to overcome the inhibition.

**Table 3.** Effect of T1 and A3-41 on D4 growth. Average *toxR* copy number for D4 cultures grown in 2216 medium mixed with T1 or A3-41 after 3- and 24-hours was determined as described in Methods. Initial cell density for strain A4 was 2 x 10^4^ CFU/ml. Results are the average of four replicates per sample.

| Addition to D4 Culture<br>(CFU/ml), final concentration | *toxR* Copy Number per ml Culture |  |
| --- | --- | --- |
|  | 3 hrs | 24 hrs |
| None | $1.6 \times 10^5 \pm 0.3 \times 10^5$ | $1.3 \times 10^9 \pm 0.5 \times 10^9$ |
| T1 @ $2 \times 10^4$ | $8.8 \times 10^5 \pm 6.5 \times 10^5$ | $1.5 \times 10^9 \pm 0.9 \times 10^9$ |
| T1 @ $2 \times 10^5$ | $8.8 \times 10^5 \pm 6.3 \times 10^5$ | $8.8 \times 10^8 \pm 1.5 \times 10^8$ |
| T1 @ $2 \times 10^6$ | $1.7 \times 10^5 \pm 0.6 \times 10^5$ | $1.2 \times 10^6 \pm 0.2 \times 10^6$ |
| A3-41 @ $2 \times 10^4$ | $7.5 \times 10^5 \pm 7.1 \times 10^5$ | $7.0 \times 10^9 \pm 2.0 \times 10^9$ |
| A3-41 @ $2 \times 10^5$ | $1.4 \times 10^6 \pm 1.0 \times 10^6$ | $3.1 \times 10^9 \pm 0.7 \times 10^9$ |
| A3-41 @ $2 \times 10^6$ | $1.9 \times 10^5 \pm 1.0 \times 10^5$ | $2.1 \times 10^9 \pm 1.6 \times 10^9$ |

A scenario that could explain the density-dependent activity observed for the co-culture experiments, which would be consistent with the production of a diffusible substance, as inferred from the overlay assay, is that T1 produces and secretes a D4-sensitive product that accumulates during late exponential and/or stationary phase, similar to other *B. subtilis* species. To test this possibility, supernatant fractions from 18 hr T1 cultures grown in 2216 broth were filtered and mixed with equal volumes of overnight D4 cultures freshly diluted into 2216, monitoring D4 growth as described in Methods. As shown in Fig. 2, while all cultures grew to a similar final density, growth of the D4 culture supplemented with the T1 cell-free supernatant was delayed for 5 to 6 hr, compared to the D4 control, while no delay was observed for the culture treated with a cell-free supernatant from an overnight D4 culture. D4 *toxR* levels for the T1 supernatant-treated culture were approx. 3×10^5^-fold lower compared to the untreated D4 culture (Table 4), 5.5 hr after inoculation. On the other hand, D4 levels in the D4 culture treated with its own supernatant declined only approx. 10-fold (Table 4), indicating that inhibition was due to a factor specifically produced by T1 and not likely due to exhaustion of nutrients in the spent medium. Inhibition was also found to be concentration-dependent since addition of T1 cell-free supernatant at 25% resulted in an approx. 2 hour lag (not shown). D4 growth was not affected when cell-free supernatants were prepared from mid-exponential T1 cultures (not shown).

**Table 4.** Effect of cell-free culture supernatant on D4 growth. Average *toxR* copy number for D4 cultures grown with supernatant fractions from 18 hr cultures (50% of the total culture volume) were determined as described in Methods. Samples were taken 5.5 hrs after inoculation. Results are the averages of four technical replicates for each sample.

| Addition to D4 culture | *toxR* copy number per ml |
| --- | --- |
| +50% 2216 medium | $5.3 \times 10^7 \pm 0.7 \times 10^7$ |
| +50% D4 supernatant | $5.0 \times 10^6 \pm 1.2 \times 10^6$ |
| +50% T1 supernatant | $1.8 \times 10^2 \pm 0.7 \times 10^2$ |
| +50% A3-41 supernatant | $1.5 \times 10^7 \pm 0.4 \times 10^7$ |

**Figure 2.**
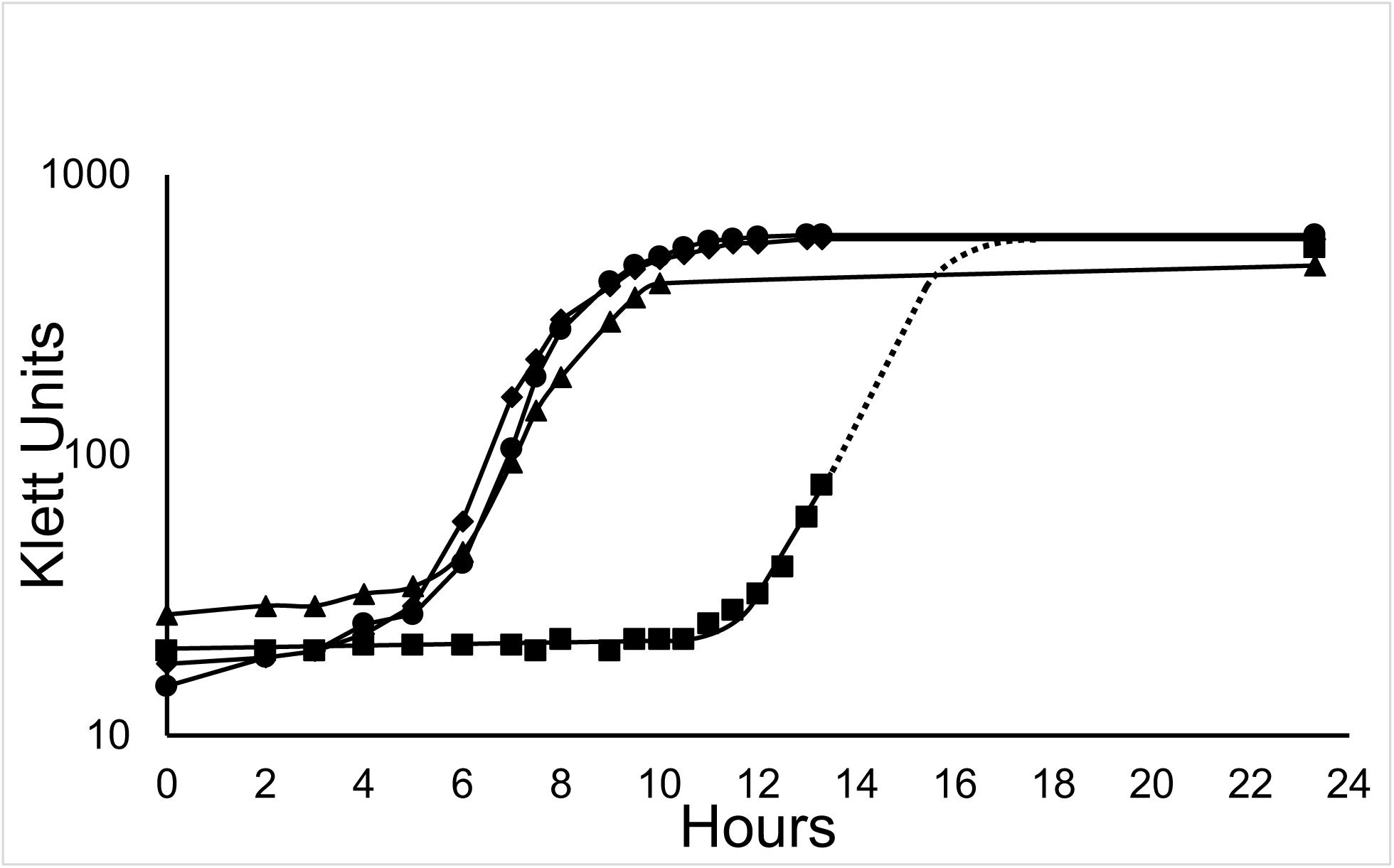
Effect of culture supernatant on D4 growth. Growth of an untreated D4 culture (●) and cultures mixed with an equal volume of cell-free supernatant fractions prepared from overnight cultures of wild-type T1 (▪), D4 (▴), and A3-41 (♦). All cultures were grown in 2216 broth. The dotted line indicates anticipated growth. Graph shows results for a representative experiment and standard deviations were <10% and were omitted for clarity.

### Identification of T1 genes involved in D4 inhibitory activity

To determine the genetic basis for the T1 inhibitory activity we generated a transposon insertion library using the *mariner*-derived *himar1* transposase as described in Methods. Over 3,000 transposon-containing mutants were screened for loss of T1 inhibitory activity by the overlay assay. Seventeen mutants were identified as having complete or partial loss of activity against D4 and insertion site locations could be identified for 16 of these mutants, which are listed in Table 5; overlay assays for mutants A2-18 (partial loss), A3-41, A11-79 and A20-86 (full loss) are shown in Fig. 3. Insertion sites for 13 mutants were found clustered in seven open reading frames within a 42.6 kb region of DNA positioned between the *pcrA-ligA* operon and *phoA* at 5’ and 3’ ends (Table 5 and Fig. 4), respectively, unique to the *inaquosorum* and *stercoris* subspecies of *B. subtilis* (18). BLASTp analysis revealed that the deduced *orfA, orfB*, and *orfC* products containing peptide synthesis, condensation, adenylation, and acyl carrier domains conserved among non-ribosomal peptide synthetases (NRPSs). Domains conserved among type I polyketide synthases (PKSs), including kedoreductase, ketoacyl synthase, and acyl transferase, were evident for *orfD, orfE* and *orfF* products. Furthermore, homologies were found to *B. subtilis* proteins involved in antimicrobial peptide and polyketide synthesis for the other open reading frame products including major facilitator superfamily (MFS) transporter (*orf2*), endopeptidase processing enzyme (*orf3*), thioesterase (*grsT*), acyl CoA dehydrogenases (*orf5* and *orf8*), and acyl carrier protein (*orf7*) (Fig. 4). The DNA sequence of this region was deposited in GenBank, accession number MT366812.

**Table 5.** Summary of transposon insertion sites in T1 mutants. Shown are the delivery vector used to create the mutant, location of the insertion site, gene and its putative product, and the nucleotide location. Accession number for the DNA sequence is MT366812. Unless otherwise indicated, nucleotide position of the insertion site is relative to the start of the *pcrA* gene, the first gene of the sequenced region. NRPS, non-ribosomal peptide synthetase; PKS, polyketide synthase.

| Mutant | Delivery Vector | Transposon Insertion Site | Gene, Putative Gene Product | NRPS-PKS ORF Location |
| --- | --- | --- | --- | --- |
| A20-86 | pDP384 | 8,534 | *orfA*, NRPS | 7,036 to 11,532 |
| S21-46 | pEP4 | 9,917 |  |  |
| A13-56 | pDP384 | 18,057 | *orf8*, Acyl CoA Dehydrogenase | 16,935 to 18,062 |
| A2-30 | pEP4 | 19,316 | *orfB*, NRPS | 18,084 to 27,179 |
| A10-67 | pDP384 | 19,316 |  |  |
| S11-14 | pEP4 | 21,936 |  |  |
| S12-29 | pEP4 | 21,936 |  |  |
| A3-41 | pDP384 | 24,490 |  |  |
| A11-79 | pDP384 | 27,466 | *orfC*, NRPS | 27,199 to 29,868 |
| S21-28 | pEP4 | 28,790 |  |  |
| A1-20 | pDP384 | 32,220 | *orfD*, PKS | 29,861 to 34,387 |
| S35-30 | pEP4 | 34,499 | *orfE*, PKS | 34,405 to 41,949 |
| A23-33 | pDP384 | 44,976 | *orfF*, PKS | 41,951 to 48,379 |
| A2-18 | pDP384 | 399* | *oppA*; ABC transporter substrate-binding protein | NA |
| S31-22 | pEP4 | 427* | *yoaZ*; putative oxidative stress response factor | NA |
| A8-11 | pDP384 | 68* | *spo0A*; sporulation master regulator | NA |

\*relative to the initiation codon of the corresponding gene in *B. subtilis* (NC\_000964.3); NA, not applicable

**Figure 3.**
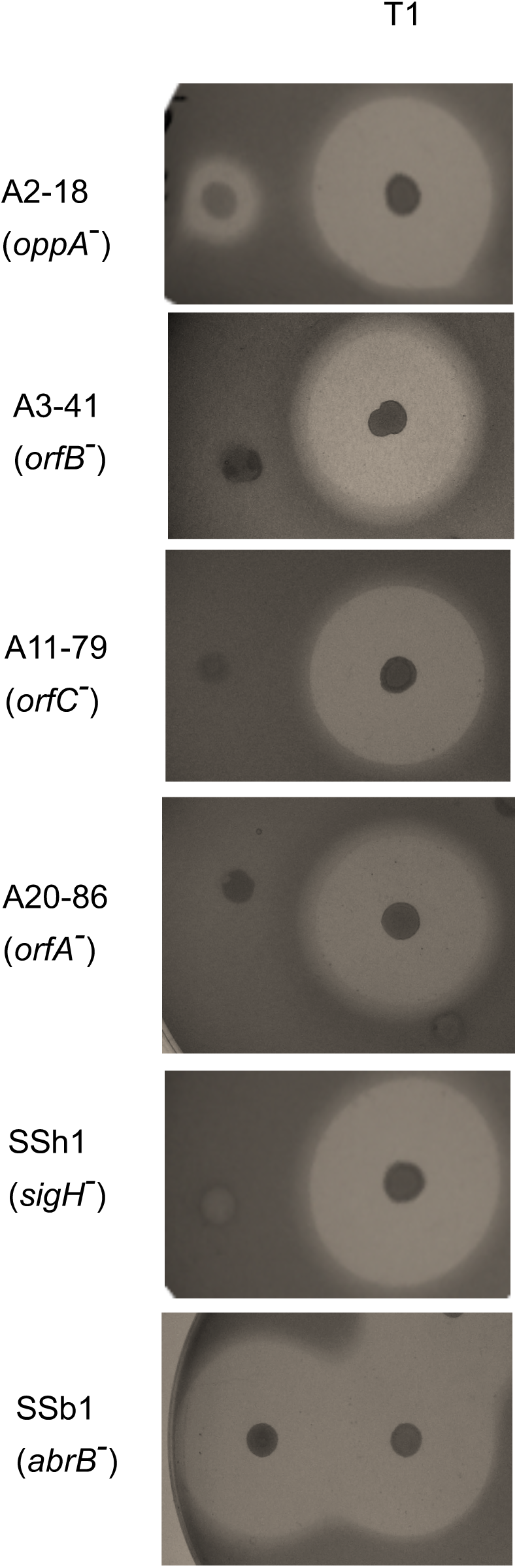
Soft-agar overlay assays for select T1 mutants (left) and wild-type T1 (right) against strain D4. Assays were done as described in Methods.

**Figure 4.**
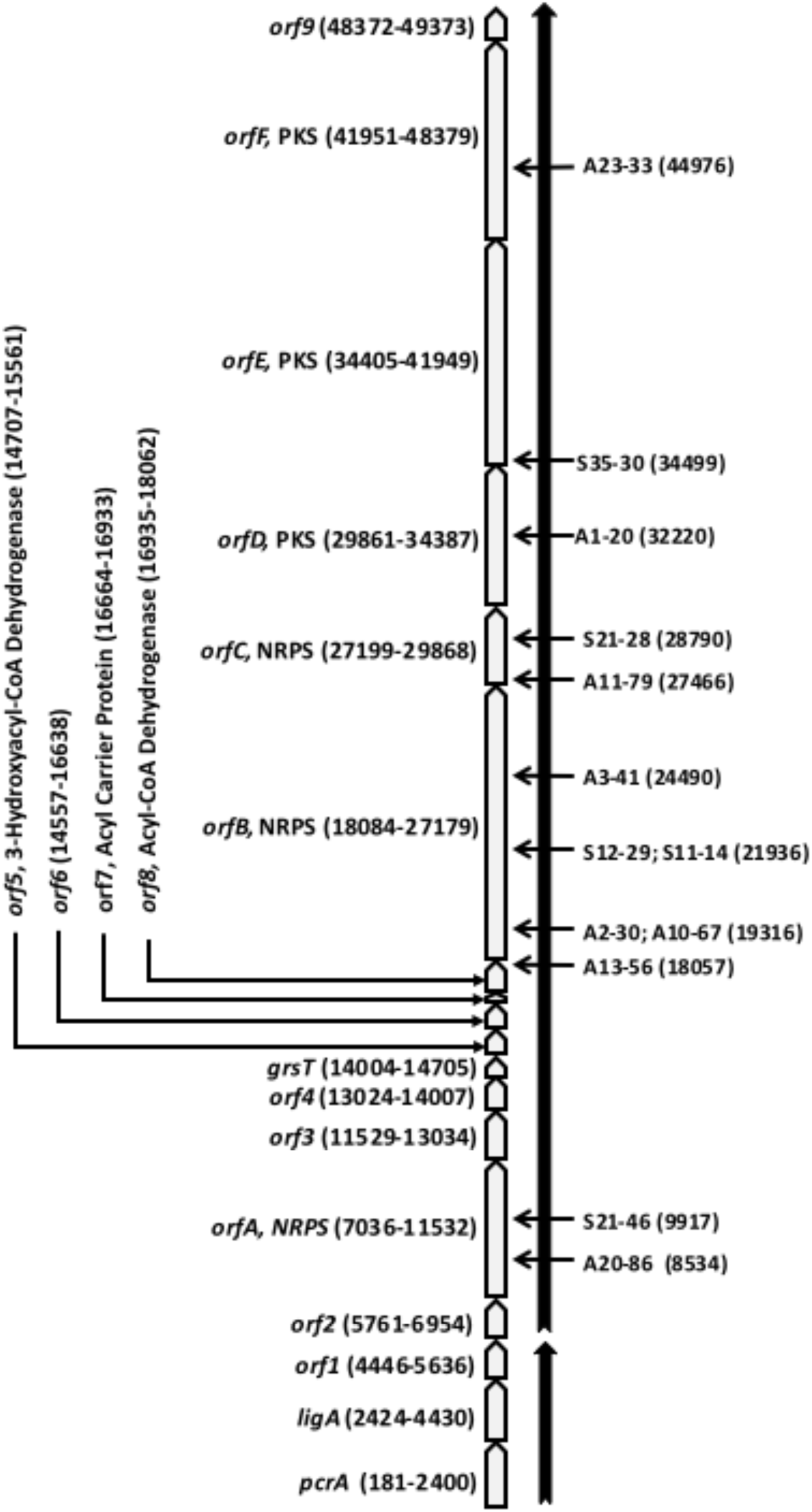
Genetic and transcription map of the NRPS-PK region of strain T1 and mutant insertion sites. See text for details.

In addition to evaluating mutant activities using the overlay assay, we examined the activity of mutant A3-41, grown in co-culture with D4 at a 1:1, 10:1 and 100:1 ratio. This mutant contains an insertion in the NRPS-like *orfB* gene, about halfway into the NRPS-PKS region (Table 5 and Fig. 4). Consistent with the overlay assay results, A3-41 did not significantly affect D4 levels at either the 3 hr or 24 hr timepoints (Table 3) when included at any ratio. Furthermore, the cell-free supernatant prepared from A3-41 did not delay D4 growth like its wild-type T1 parent (Fig. 2) and D4 levels at 5.5 hr after inoculation in the presence of the A3-41 cell-free supernatant were comparable to the control D4 culture (Table 4). At 24 hr, A3-41 levels (*amyE* copy number) were not significantly different and were similar to T1g levels, varying between 2.4×10^8^ ± 0.5 x10^8^ copies/ml for the 1:1 culture to 4.7×10^8^ ± 1.2×10^8^ copies/ml for the 100:1 culture. Thus, loss of inhibitory activity was directly due to elimination of the *orfB* product and consistent with a role for the NRPS-PKS region in producing the anti-D4 activity.

Mutants with insertions outside of the NRPS-PKS cluster were also found, including within *spo0A* (mutant A8-11, Table 5), the master transcriptional regulator required for activating early sporulation and early stationary phase genes (25), and *oppA* (mutant A2-18, Table 5). Also known as *spo0K, oppA* is the first gene of a five-gene operon responsible for synthesis of the oligopeptide-binding protein an ABC-transporter that plays a role in a variety of stationary phase activities, including initiation of sporulation and competence development (26). In both cases, gene disruption resulted in partial loss of D4 inhibitory activity (Fig. 3 and not shown). Mutant S31-22 (Table 5), which was also defective in D4 inhibitory activity, contained an insertion in *yoaZ* (BSU17890), a putative oxidative stress response factor whose function is unclear (27).

### Transcription mapping of the NRPS-PKS region

RNA extracted from mid-exponential and early stationary phase cultures of T1 grown in 2216 broth was examined by RT-PCR, as described in Methods. Primers were designed to target amino and carboxyl termini of consecutive predicted ORFs (Table 2). Regions spanning all 17 predicted ORFs in the NRPS-PKS region were tested as well as the adjacent upstream *pcrA* operon and downstream *phoA* gene. Amplification of cDNA generated from both mid-exponential (not shown) and stationary-phase (Fig. 5) RNA was observed for the entire NRPS-PKS region spanning from *orf2* to *orf9*, indicating that the entire 42.6 kb region encompassed a single transcription unit. No amplification products were observed for the *orf1-orf2* and *orf9-phoA* regions (Fig. 5), signifying the absence of a transcript spanning these genes and demarcating the 5’ and 3’ ends of the NRPS-PKS region transcriptional unit at *orf2* and *orf9*, respectively. The absence of a product between *orf9* and *phoA* was expected since *phoA* is expressed in a direction opposite of the NRPS-PKS region. Both *orf9* and *phoA* stop codons are followed by sequences that could form 25 and 27 b mRNA hairpin structures (ΔG= -17 kcal/mol for both, determined using the Vienna RNA Websuite (28)) with U-rich ends, which likely act as a transcription terminators. Similarly, *pcrA, ligA* and *orf1* are part of a four-gene operon—*pcrA* is preceded by *pcrB* and *orf1* (referred to as *yerH* by Petit et al. (29))— with a transcription terminator found after *orf1* (see below), so co-transcription of *orf1-orf2* was not anticipated.

**Figure 5.**
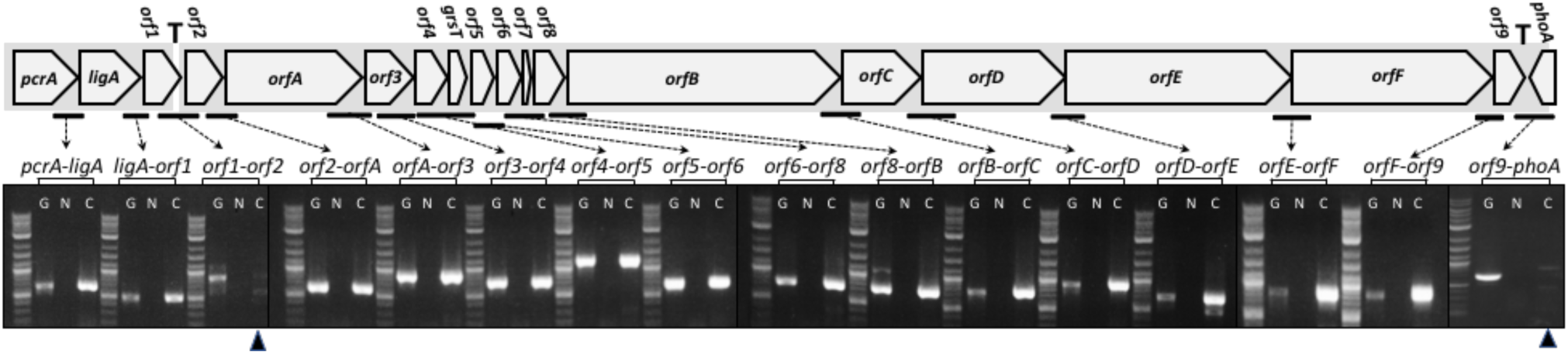
Transcription mapping of the NRPS-PK region. Predicted open reading frames and gel electrophoresis of RT-PCR products generated using T1 RNA isolated from stationary phase cultures (T_1_ to T_2_). Primer pairs targeting open reading frame termini in the NRPS-PK region as well as upstream and downstream genes, as indicated, are described in Methods and Table 2. Lanes with a G are PCR reactions containing genomic DNA (-RT, -DNase); N, PCR with RNA and DNase but without RT, and; C, RT-PCR of RNA after DNase treatment. The absence of RT product for *orf1-orf2* and *orf9-phoA* regions is indicated by triangles (▴); potential transcription terminators are indicated by a “T”.

### Control of NRPS-PKS gene expression

Several transposon mutants were obtained using plasmid pEP4, which, upon insertion, resulted in the creation of a transcriptional fusion with *lacZ*, thereby enabling the analysis of gene expression during growth and stationary phase. ß-Galactosidase levels were determined for four mutants, A20-86, A3-41, A11-79 and A1-20, containing fusions with *orfA, orfB, orfC* and *orfD*, respectively (Table 1). These mutants were grown in 2216 broth and harvested at mid-exponential (T_-1_), at the transition into stationary phase (T_0_), and 1-2 hrs into stationary phase (T_1_) and results are shown in Table 6. During mid-exponential growth, ß-galactosidase levels were similar for all four strains and did not fluctuate significantly compared to background levels obtained for wild-type T1g, which does not possess *lacZ*. Once cultures entered stationary phase, *lacZ* expression increased between 17- to 42-fold compared to their T_-1_ basal levels and were elevated an additional 50% 1-2 hrs into stationary phase. Thus, expression of NRPS-PKS genes are linked to stationary phase processes, consistent with the isolation of mutants in stationary phase regulator genes.

**Table 6.** ß-Galactosidase activities of T1 strains at different stages of growth.

| Strain | Relevant genotype | β-Galactosidase Activity (nmoles/min/OD<sub>600</sub>) | | |
| --- | --- | --- | --- | --- |
|  |  | Growth Stage* |  |  |
|  |  | T<sub>-1</sub> | T<sub>0</sub> | T<sub>1</sub> |
| T1g |  | 167±29 | 108±39 | 206±79 |
| A20-86 | *orfA::lacZ* | 140±51 | 5805±158 | 8660±610 |
| A3-41 | *orfB::lacZ* | 224±53 | 3898±35 | 5941±180 |
| A11-79 | *orfC::lacZ* | 180±59 | 4670±1060 | 7012±650 |
| A1-20 | *orfD::lacZ* | 115±24 | 3905±714 | 5440±124 |
| A20-86-A | *orfA::lacZ; abrB::erm* | 7288±498 | 5888±296 | 7240±260 |

Strains were grown in 2216 broth and $\beta$ -galactosidase levels were measured as described in
Methods. \*T<sub>-1</sub>, OD<sub>600</sub> ~1; T<sub>0</sub>, OD<sub>600</sub> ~3; T<sub>1</sub>, OD<sub>600</sub> 4-5

### The *orf1-orf2* intergenic region

Examination of the 117 bp region between *orf1* and *orf2* revealed several potential promoter and regulatory sequences. We identified a sequence adjacent to the predicted *orf1* stop codon having the capacity to form a 30 b mRNA hairpin structure (ΔG= -21 kcal/mol) followed by several consecutive Us, which likely serve as a transcription terminator (Fig. 6). Sequences between 36 and 57 bp upstream from the *orf2* initiation codon (Fig. 6) were similar to σ^A^ (30) and σ^H^ (31) consensus sequences and may be used for *orf2* transcription initiation. The loss of D4 inhibiting activity by strain SSh1 (Fig. 3), which contains an insertion in *sigH*, the σ^H^ structural gene, supports involvement of σ^H^ in NRPS-PKS gene expression. Similarly, the partial loss of inhibiting activity observed for *spo0A* mutant A8-11 in the overlay assay also indicates a role for Spo0A in activating NRPS-PKS expression. A putative *spo0A* binding site was identified 71 bp upstream of the σ^A^ and σ^H^ consensus sequences, overlapping the last two *orf1* codons; this sequence includes internal G and C residues critical for Spo0A binding (25) (Fig. 6). The location of this site could allow for Spo0A-dependent activation of either σ^H^ or σ^A^ polymerases.

**Figure 6.**
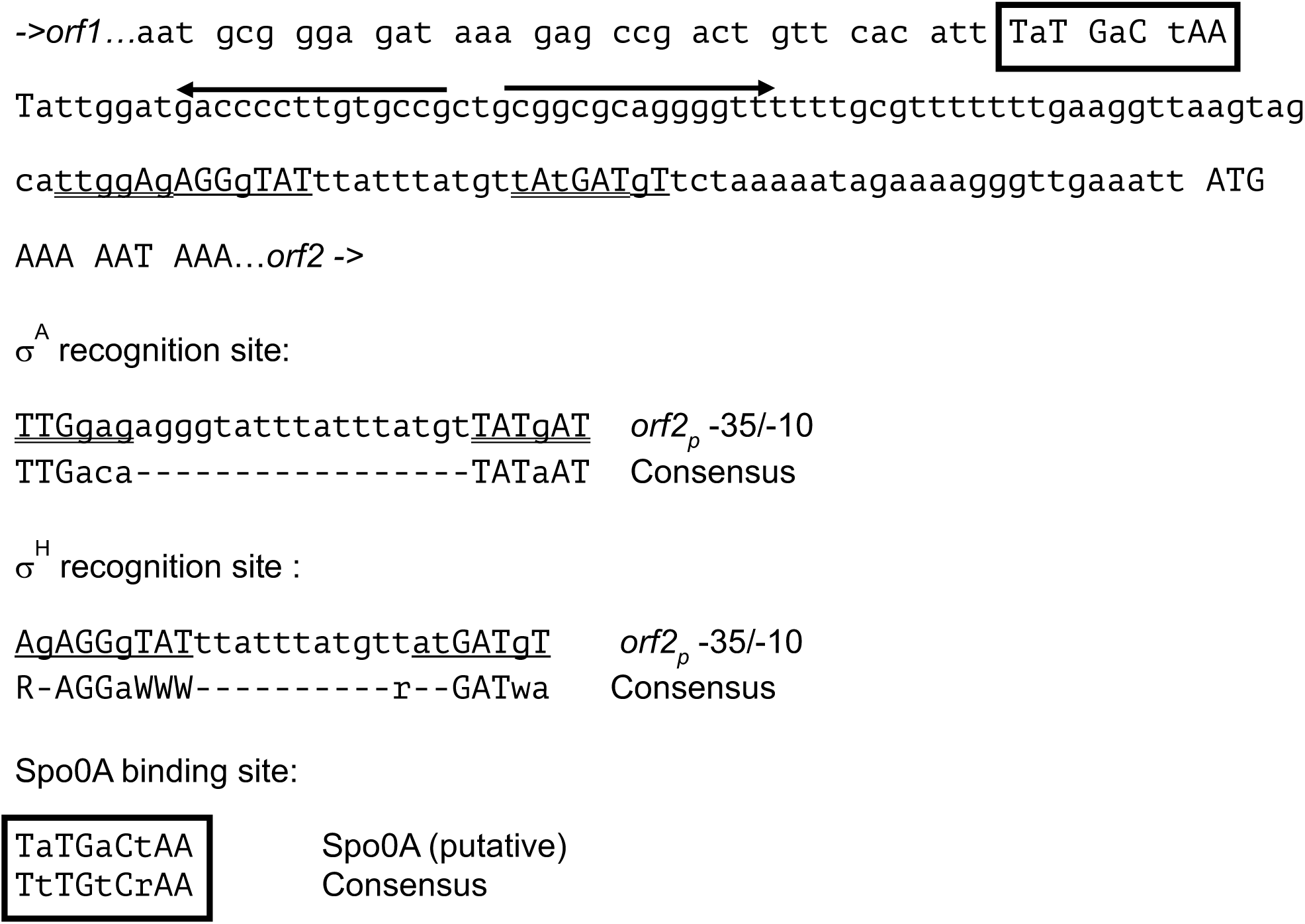
The *orf2* promoter region. Putative regulatory and promoter features for the DNA sequence between the last 13 codons and first four codons of *orf1* and *orf2* open reading frames, respectively, are shown. Also shown are the consensus sequences and recognition sites for σ^A^ (double underlined), σ^H^ (single underlined), and Spo0A (boxed). Bases that agree with consensus sequences are capitalized.

### Involvement of AbrB on NRPS-PKS expression

AbrB is a key transitional phase regulator controlling expression of more than 100 stationary phase genes including those involved in antibiotic production (32). To examine a role for AbrB in NRPS-PKS expression, we constructed *abrB* mutant SSb1(*abrB*::*erm)*. Like its parent T1, SSb1 was found to inhibit D4 growth using the overlay assay (Fig. 3). A cell-free supernatant prepared from a stationary phase SSb1 culture delayed D4 growth approx. 22 hr, four-fold longer than the delay observed for the T1 supernatant (Fig. 7). Furthermore, ß-galactosidase activity of *abrB* mutant A20-86A (*orfA::lacZ*) was 52-fold higher during mid-exponential (T_-1_) than strain A20-86, its *abrB*^*+*^ parent, and comparable to the elevated levels observed for transition (T_0_) and stationary phases (T_1_) (Table 6). Therefore, AbrB plays a role in controlling NRPS-PKS expression.

**Figure 7.**
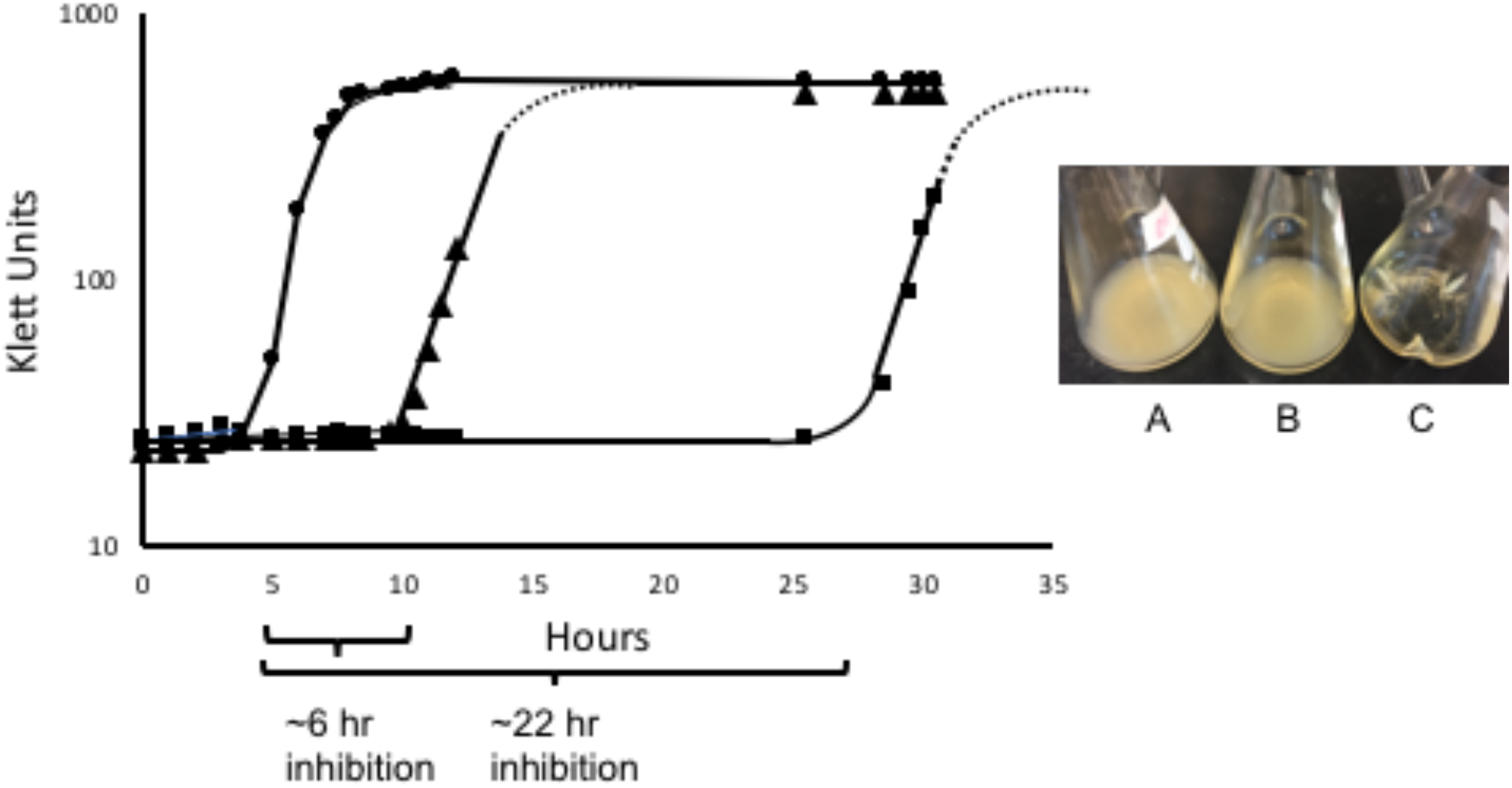
Deletion of *abrB* results in enhanced anti-D4 activity. Growth of an untreated D4 culture (●) and cultures mixed with an equal volume of cell-free supernatant fractions prepared from overnight cultures of wild-type T1 (▴) or SSb1(*abrB*^*-*^) (▪) as described in Methods. Dotted lines indicate anticipated growth. Graph shows results for a representative experiment and standard deviations were <10% and were ommitted for clarity. Inset: photograph of 26-hour cultures; (A) D4 alone, (B) D4 with cell-free superatants from T1, (C) D4 with cell-free supernatant of SSb1.

## DISCUSSION

Our preliminary studies showed that *B. subtilis* strain T1 inhibited growth of VirAP toxin-producing *V. parahaemolyticus* (Avery, Hise, and Schreier, unpublished). Analysis of the T1 genome revealed that it is a member of the *inaquosorum* subspecies of *B. subtilis* (Schreier, unpublished), which is distinguishable from *subtilis* and *spizizinii* subspecies by a 42.6 kb DNA region having the capacity to encode NRPSs and PKSs (18). In the present study we used transposon mutagenesis to determine that this region is responsible for producing the inhibitory activity and results from overlay assays and cell-free culture supernatant experiments indicates that this activity is a secreted stationary phase product whose synthesis is under control of key stationary phase regulators. To our knowledge, this is the first study that examines expression and control of an antibacterial compound from the NRPS-PKS region unique to the *inaquosorum* and *stercoris* subspecies of *B. subtilis* (18).

Synthesis and secretion of antimicrobial compounds by members of the genus *Bacillus* during the transition to stationary and early stationary phase is one strategy used by this group of bacteria to compete with other organisms for reduced resources as a prelude to sporulation (33). The elevated expression observed for NRPS-PKS genes by T1 during these periods (Table 6) is consistent with this strategy and, with the the low or negligible expression during growth, can explain results for the D4 and T1 co-cuture experiments. During exponential growth (i.e., 3 hr after inoculation), the inability to inhibit D4 growth by any of the T1 treatments is likely the consequence of low level inhibitor production during this period. Throughout the subsequent 21 hrs, however, the entry of T1 cultures into late exponential and stationary phase resulted in as much as a 62-fold increase in NRPS-PKS gene expression (e.g., for strain A20-86, Table 6) with the accompaying synthesis and accumulation of inhibitor. This resulted in a 700- to 1200-fold decrease in D4 levels observed for growth in the presence of 100-fold excess T1 (Table 3). On the other hand, the absence of inhibition observed for D4 in co-cultures of T1 at 1:1 and 10:1 ratios could be explained by differences in generation times between D4 and T1 in 2216 broth, which are 40 and 60 min, respectively. For these cultures, D4 likely outgrew and attained stationary before T1 accumulated sufficient inhibitor required to influence D4 growth. Furthermore, at the 1:1 and 10:1 ratios, D4 may be able to mitigate the effect of inhibitor, as was observed by its ability to resume growth after treatment with cell-free T1 culture supernatants (see below).

Evidence that D4 was sensitive to a product of the NRPS-PKS region was obtained from overlay, co-culture, and cell-free supernatant experiments, since inhibition did not occur using mutants containing insertions within NRPS-PKS region genes. Moreover, the inability of cell-free supernatants from *orfB* mutant A3-41 to inhibit D4 growth indicated that the effect was directly due to *orfB* and downstream genes and not a consequence of a toxic stationary phase byproducts, e.g., volatile organic or inorganic compounds (34), or nutrient depletion of the spent medium. While *B. subtilis* subsp. *inaquosorum* is capable of synthesizing lipopeptides bacillomycin F and fengycin (18), growth of *V. parahaemolyticus* strains A3, D4, and ATCC 17802 occurred in the presence of both T1 NRPS-PKS mutants as well as strain SMY, a *B. subtilis* subsp. *subtilis* that also produces these secondary products, arguing that neither of these nonribosomal peptides are involved in T1 inhibitory activity.

Transcript mapping indicated that the NRPS-PKS region encodes one polycistronic message extending across all 15 genes forming an operon, although our analyses cannot rule out the possibility that transcription of a subset of these genes might occur from internal promoters. The absence of a transcript spanning *orf1* and *orf2* indicated that initiation likely occurs from the first gene of the operon, *orf2*, and requires σ^H^ for activity (based on results for the overlay assay of *sigH* mutant SSh1, Fig. 3), which may bind to sequences upstream of *orf2* (Fig. 6). Recent RT-PCR studies have detected NRPS-PKS mRNA in stationary phase cultures of strain SSh1 (Ruzbarsky, unpublished), suggesting the involvement of another polymerase for transcription. One candidate is σ^A^ since consensus σ^A^ promoter sequences were found to overlap the putative σ^H^ promoter (Fig. 6), and would provide a mechanism for NRPS-PKS expression during exponential growth. Promoters transcribed using both sigma factors under different physiological conditions in *B. subtilis* have been noted (31, 35).

Similar to many *B. subtilis* stationary phase products and processes (33), the NRPS-PKS genes of T1 were found to be under control of *oppA, spo0A* and *abrB*, genes encoding transitional and stationary phase regulators. Disruption of *oppA*— mutant A2-18—the first gene of the *opp* operon, resulted in loss of inhibitory activity. This operon encodes an oligopepetide permease that functions as a receptor in signaling (26, 36) and impairment of *oppA* results in loss of bacilysin production (37), a nonribosomally synthesized antimicrobial. The *opp* operon is linked to the ComA competence response regulator and CSF (PhrC), the competence and sporulation factor, which also participates in *B. subtilis* quorum-sensing (36). When CSF accumulates extracellulary due to high cell density, production of bacilysin is stimulated (38). Any involvement of the Com system or CSF in control of NRPS-PKS expression through *oppA* is yet to be determined.

Isolation of a mutant with an insertion in *spo0A*—A8-11—having decreased inhbitory activity indicated a role for this regulator in NRPS-PKS control. The *spo0A* gene product, Spo0A, is a master transcriptional regulator of early stationary phase processes, including NRPS and PKS gene activation, and development of spores (39). In its phosphorylated state, Spo0A activates transcription initiation at both σ^A^ and σ^H^ promoters and is responsible for indirectly activating *sigH* transcription by repressing AbrB, the global transition state regulator (33, 39). While we did not address the nature of Spo0A participation on NRPS-PKS expression, we identified a putative Spo0A binding site upstream of *orf2* that could be used to activate σ^H^- and, possibly, σ^A^-dependent transcription. Whether Spo0A is directly involved in activating NRPS-PKS transcription or indirectly by elevating σ^H^ levels remains to be established.

Like Spo0A, AbrB is essential for controlling NRPS-PKS expression as *orfA* expression in *abrB* mutant A20-86-A was found to be 52-fold elevated during exponential growth compared to its parent A20-86 (*abrB*^+^) strain (Table 6). Elevated activity was also found for cell-free supernatant fractions prepared from the *abrB* mutant, SSb1 (Fig. 7). Involvement of AbrB in controlling NRPS and PKS genes in *Bacillus* spp. is well documented, and can act by binding either directly to promoter sequences, interfering with transcription initiation, or indirectly through repression of *sigH* (33). While the A/T-rich character of the *orf2* promoter is typical of AbrB binding sites, TGGNA and TNCCA motifs associated with AbrB (40) are absent.

How is *V. parahaemolyuticus* affected by the T1 inhibitor? The mode of action of many *Bacillus* spp. NRPS- and PKS-derived anitmicrobials is through membrane perturbation or depolarization (15), e.g., fengycin (41) and iturin A (42), blockage of peptidoglycan biosynthesis, e.g., bacilysin (34), or selective inhibition of protein synthesis, e.g., difficidin (34). Regardless of mechanism, both co-culture and cell-free supernatant experiments demonstrated that D4 was capable of overcoming the T1 NRPS-PKS inhibitory activity, with growth levels reaching those obtained for untreated cultures. Along with D4, the bacteriostatic nature of the T1 inhibitor was also observed for A3 and ATCC 17802 strains, which produced small colonies in overlay assay clearance zones after long-term incubation (Schreier, unpublished). Isolates obtained from these zones continued to remain sensitive to T1 (Schreier, unpublished), suggesting an ability to either degrade the inhibitor—*Vibrio* spp. secrete several classes of proteases, some of which may have activities against lipopeptides (43)—or modify their membrane components to be less sensitive to inhibitor activity as part of an induced stress response (44). The increased sensitivity of A3 to T1 inhibition compared to D4 and ATCC 17802 in the overlay assay might be explained by increased activities for either of these processes. While little is known about stability and persistance of NRPS and PKS products in the environment, it is also possible that factors other than enzymatic degradation, e.g., pH and media components, may decrease their efficacy over time (45).

The use of *Bacillus* spp. as biological control agents has received a substantial amount of attention due to their antimicrobial characteristics and safety (13, 46) along with their spore-forming capability, which is advantageous for long-term storage (47). *B. subtilis* strains have been effective in controlling disease outbreaks due to *Vibrio* pathogens in a variety of aquaculture species, including shrimp (48-50), in addition to providing probiotic benefits (13). The ability of *B. subtilis* strain T1 to inhibit growth of AHPND *Vibrio* spp. suggested its potential for use as a tool in the prevention and treatment of AHPND in shrimp aquaculture. Understanding the genetic basis of the inhibitory activity and its regulatory mechanisms allows for the development of T1 strains having desirable properties such as enhanced NRPS-PKS expression during growth that was acquired by disabling *abrB*. Such strains could be used in aquaculture applications to complement other strains having different antimicrobial activities. Our preliminary studies have shown that daily addition of freshly prepared cell-free SSb1 supernatants to D4 cultures resulted in continuous cessation of D4 growth over the course of a 72 hour treatment (Avery, Ruzbarsky and Schreier, unpublished). Thus, supplementing feed with *abrB* mutant SSb1 might provide system water and animal microbiomes with a constant source of inhibitory activity, bypassing stability or degradation issues and restricting the growth of pathogen before they reach virulent levels. Studies aimed at evaluating T1 and its derivatives to protect against AHPND *Vibrio* in shrimp aquaculture systems are ongoing.

## AUTHOR CONTRIBUTIONS

SA, SR and HS conceived the project and designed the research. HS supervised the study. SA, SR, AH and HS contributed to the investigation, methodology and data analysis. SA, SR and HS prepared figures, wrote, reviewed and edited the paper. All authors reviewed and approved the final manuscript.

## ACKNOWLEDGEMENTS

This works work was supported, in part, by grants from Epicore Networks (U.S.A.) Inc. and G. Unber Vetlesen Foundation. The authors thank Julie Wolf, Dr. Eric Schott and Dr. Russell Jerusik for helpful comments and Sabeena Nazar, Bioanalytical Services Lab, Institute of Marine and Environmental Technology, for DNA seqeuncing.

